# Opposing behavioral roles for a single gene in a species with a supergene polymorphism

**DOI:** 10.64898/2026.04.15.718755

**Authors:** Mackenzie R. Prichard, Isabel Fraccaroli, Erik N. K. Iverson, Donna L. Maney

## Abstract

Behavioral trade-offs, such as between territorial aggression and parental care, require coordination among neural and genetic mechanisms that influence opposing phenotypes. In some systems, including the white-throated sparrow, this coordination has been proposed to be mediated in part by chromosomal inversions that bind together multiple genes associated with alternative strategies. Here, we demonstrate that opposing components of a life-history trade-off can be linked, via alternative alleles inside a chromosomal rearrangement, to a single pleiotropic gene. In white-throated sparrows, a species with polymorphic aggressive and parental behavior, the gene encoding the neuromodulator vasoactive intestinal peptide (VIP) lies within an inversion-based supergene. Working with a free-living population, we found that *VIP* expression in a pro-aggression cell population in the anterior hypothalamus positively predicted territorial singing, whereas expression in a pro-parental cell population in the infundibular nucleus positively predicted parental provisioning. Expression in these two regions was negatively correlated, consistent with opposing behavioral roles. Allele-specific analyses revealed stronger expression bias toward the supergene-associated allele in the anterior hypothalamus than in the infundibular nucleus. These findings suggest that chromosomal inversions can support opposing behavioral phenotypes not only by linking multiple genes, but also by driving regulatory divergence of individual genes that organize behavioral trade-offs.

## Introduction

Behavioral trade-offs are a central theme in evolutionary biology. In many vertebrates, for example, aggression and parental care are typically conceptualized as opposing investments: individuals that devote more effort to territorial defense are expected to invest less in offspring, and vice versa (Magrath & Komdeur, 2003; Trivers, 1972). In behavioral neuroendocrinology, aggression and parenting are typically treated as functionally and mechanistically separable, each supported by its own neural and genetic architecture. When behaviors are coupled as components of a trade-off, however, selection may act on combinations of alleles influencing both traits, potentially favoring their co-adaptation at the genetic level.

Chromosomal inversions represent a well-established genomic mechanism for promoting and maintaining co-adaptation of alleles that influence multiple, related traits. By suppressing recombination across large genomic regions, inversions allow sets of alleles to be inherited together as a single haplotype. According to Dobzhansky (1950), such arrangements can be adaptive when they capture combinations of alleles that function well together—for example, variants in a hormone and its receptor—thereby creating and preserving co-adapted gene complexes. In this way, “supergenes” have sometimes been invoked to explain the evolution of alternative life histories and social strategies in which suites of traits, including those representing trade-offs, are inherited as integrated packages (Schwander et al., 2014; Thompson & Jiggins, 2014).

The white-throated sparrow (*Zonotrichia albicollis*) provides a striking example of this phenomenon. In this species, a supergene comprising a series of chromosomal inversions has resulted in the differentiation of two discrete plumage morphs with alternative behavioral phenotypes (Campagna, 2016; Maney, 2008). Individuals of the ‘white-stripe’ (WS) morph exhibit heightened territorial aggression and reduced parental provisioning, whereas birds of the ‘tan-striped’ (TS) morph show lower aggression and greater parental care (Figure 1a; Tuttle, 2003). These opposing behaviors are reliably associated with the supergene, which is present only in the WS birds.

**Figure 1.**
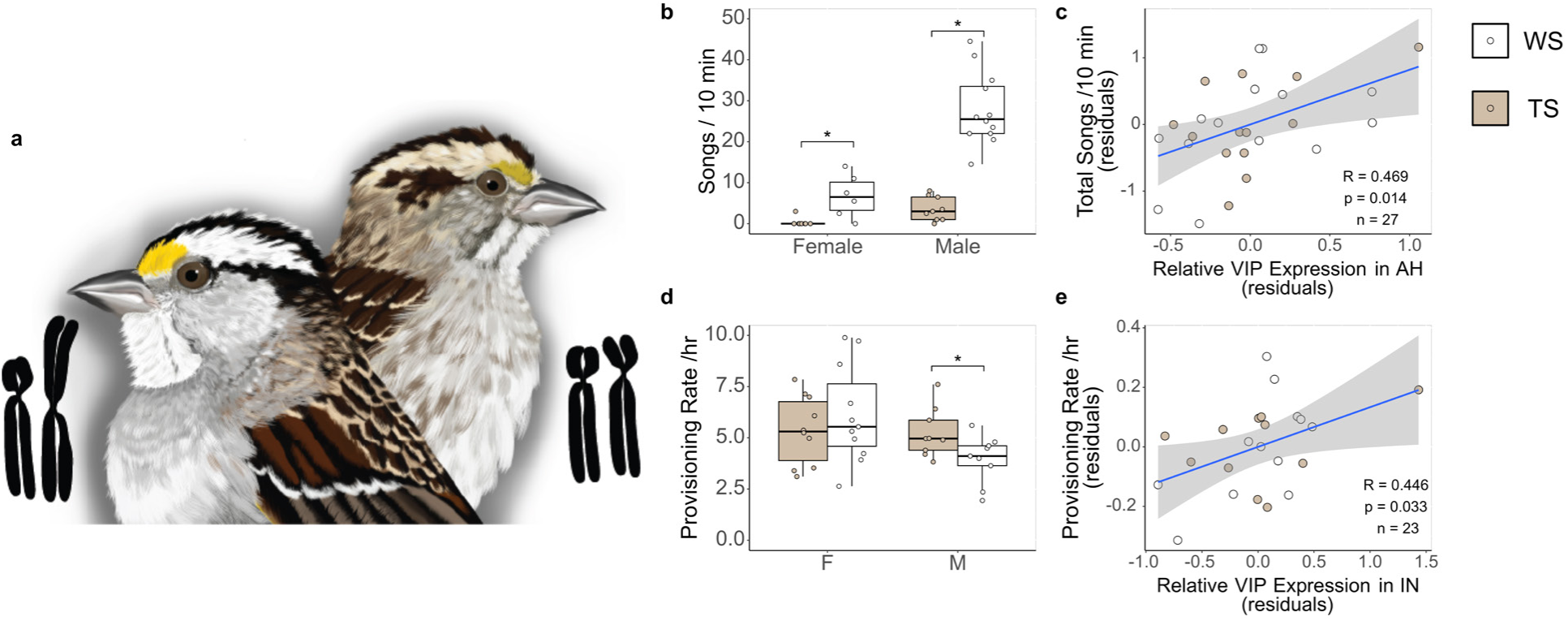
Alternative phenotypes in white-throated sparrows and associated *VIP* expression in the hypothalamus. **a,** White-striped (WS) birds (left) of both sexes have bolder, brighter coloration and carry at least one copy of ZAL2^m^, which contains a large, inversion-based supergene. Tan-striped (TS) birds (right) have more muted coloration and are homozygous for the standard chromosome, ZAL2. Illustration by M. R. Prichard. **b,** The morphs differed in territorial song rate in response to a simulated territorial intrusion**. c,** Territorial singing was positively predicted by *VIP* expression in the anterior hypothalamus (AH). **d,** Provisioning rate differed between morphs in males**. e,** Nestling provisioning rate was positively predicted by *VIP* expression in the infundibular nucleus (IN).

Under a classical framework, the supergene in white-throated sparrows is hypothesized to operate through the accumulation of distinct, co-adapted alleles, resulting in potentially large gene complexes that together support the expression of the high-aggression, low-parental behavioral strategy. Here, we consider a complementary (but not mutually exclusive) possibility: that the balance between aggression and parenting is influenced by evolutionary change in a single pleiotropic gene. We focus on vasoactive intestinal peptide (VIP), a conserved neuromodulator implicated broadly in the regulation of social behavior in vertebrates. In other songbirds, inhibition of *VIP* expression in the anterior hypothalamus (AH) reduced aggression, demonstrating a causal role for the neuromodulator in that behavior (Goodson et al., 2012a,b). In addition, VIP synthesized in the infundibular nucleus (IN) stimulates prolactin release via the portal vasculature (Maney et al., 1999; Vleck & Patrick, 1999); prolactin is widely implicated in parental behavior across vertebrates (reviewed by Smiley, 2019). Thus, the same neuromodulatory gene regulates both sides of the aggression–parenting trade-off, operating in distinct neuroanatomical contexts (see also Kingsbury, 2015; Kingsbury & Wilson, 2016). In white-throated sparrows, the protein-coding sequences of the two alleles (*i.e.*, the supergene and the standard allele, Figure 1a) are similar, with no nonsynonymous changes between them (Sun et al., 2018). Instead, there is potentially important regulatory and epigenetic variation that likely leads to differences in allele-specific expression (Prichard et al., 2022). Allele-specific elements within this gene therefore have the potential to interact with local trans-acting factors that are specific to a brain region, which would provide an efficient mechanism by which two alternative strategies in a behavioral trade-off could become genetically coordinated.

In this study, we measured territorial and parental behaviors in female and male white-throated sparrows breeding in their natural habitat. In the same sample of individuals, we quantified *VIP* mRNA expression –both total abundance and the relative expression of the two alleles—in the AH and IN. Using these measures, we sought to determine the relationships between haplotype-linked regulatory variation in *VIP* and the opposing life-history strategies associated with the supergene polymorphism in this species. These relationships could inform our understanding of how life-history trade-offs are organized at the level of gene regulation.

## Results

### *VIP* expression predicted territorial singing and parental provisioning

To test the association between *VIP* expression and aggression, we measured territorial behavior in a sample of free-living females and males that had paired and established territories but had not yet begun incubation (“pre-parental cohort,” Table S1). We conducted simulated territorial intrusions (STI) in which two observers placed a caged, male conspecific decoy and a speaker playing conspecific song within the territory of a focal pair and recorded the behavioral responses of that pair (Horton et al., 2014a). We focused our analyses on territorial singing because it is a predominate component of territorial aggression in this species and was previously shown to differ robustly between the morphs (*e.g.*, Horton et al., 2014a; Kopachena & Falls, 1993). As expected, we found that WS birds sang more often than TS birds in response to STI (F_(1,41)_ = 53.61, p < 0.001; Figure 1b, Table S2).

We collected brain tissue from all focal animals 1-2 days following STI. Using RT-qPCR, we found a significant positive correlation between *VIP* expression in the AH and territorial singing; the higher the level of *VIP* expression in this region, the more frequently the individual sang in response to STI (R = 0.469, 95% CI [0.108, 0.720], p = 0.014; Figure 1c). The correlation between *VIP* expression and singing was not significant in the IN, a region in which the VIP cell population is not associated with aggression (R = −0.104, 95%CI [-0.512, 0.343], p = 0.655; Figure S1), and this relationship was significantly weaker than in the AH (Fisher’s Z: p = 0.045). Thus, *VIP* expression in the AH, but not the IN, significantly predicted territorial singing.

To test the association between *VIP* expression and parental behavior, we quantified parental provisioning by recording videos of free-living females and males feeding nestlings that were six days post-hatch (“parental cohort,” Table S2). TS males provisioned their nestlings more often than did WS males (t = 2.166, p = 0.038; Figure 1d). As others have reported (*e.g.*, Horton & Holberton, 2010; Horton et al., 2014a), we did not detect a morph difference in females (t = −1.077, p = 0.290; Figure 1d).

*VIP* expression in the IN was positively correlated with provisioning rate; when controlling for both morph and sex, we found that the higher the *VIP* expression in the IN, the more the individual fed its nestlings (R = 0.446, 95%CI [0.041, 0.725], p = 0.033; Figure 1e). In the AH, *VIP* expression was not significantly correlated with provisioning rate (R = −0.088, 95%CI [-0.501, 0.357], p = 0.704; Figure S2), and the relationship between *VIP* expression and provisioning trended stronger in the IN than in the AH (Fisher’s Z: p = 0.080). Thus, *VIP* expression in the IN, but not the AH, significantly predicted parental provisioning.

### Differential *VIP* expression was associated with life-history strategy

The associations between *VIP* expression and behavior (Figure 1c, e; see also Horton et al., 2020) support our hypothesis that the neuromodulator plays distinct roles in behavior depending on the brain region (AH or IN). Our analysis showed morph differences in *VIP* expression in both regions, and the direction of this difference depended on the region (morph x region interaction, F(1,119) = 12.92, p = <0.001; Figure 2a, Table S4). Relative to the TS birds, the WS birds had higher levels of *VIP* expression in the AH (t = −2.856, p = 0.005; Figure 2a) and lower levels of *VIP* expression in the IN (t = 2.108, p = 0.037; Figure 2a), consistent with a pro-aggression role for the population of VIP cells in AH and a pro-parental role for the population in IN. Further, *VIP* expression in the AH was negatively correlated with *VIP* expression in the IN (R = −0.318, 95%CI [-0.542, −0.052], p = 0.020; Figure 2b). This negative relationship suggests oppositional regulation of *VIP* expression in the AH and IN, consistent with a role for this neuromodulator in a behavioral trade-off.

**Figure 2.**
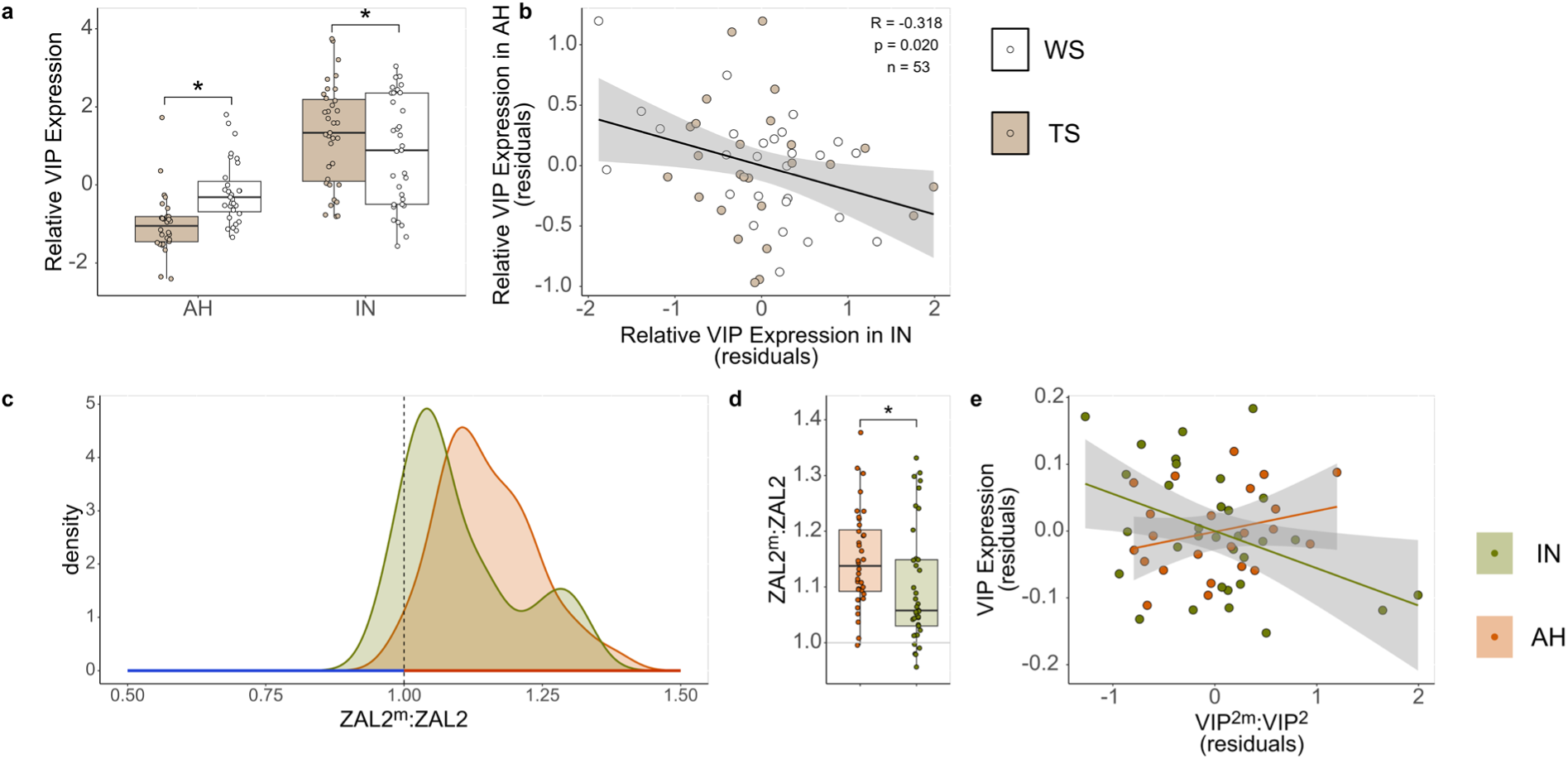
Oppositional regulation of *VIP* expression in the anterior hypothalamus and infundibular nucleus is associated with life history strategy. **a,** *VIP* expression in the anterior hypothalamus (AH) was greater in individuals of the white-striped (WS) morph (white) than the tan-striped (TS) morph (tan). The morph difference in the infundibular nucleus of the hypothalamus (IN) went in the opposite direction. **b,** *VIP* expression in the AH and the IN were negatively associated in both morphs. **c,** In both the anterior hypothalamus (AH, orange) and the infundibular nucleus of the hypothalamus (IN, green), *VIP*-2^m^ was overexpressed relative to *VIP*-2. The dashed line indicates equal expression of the alleles, greater expression of *VIP*-2^m^ relative to *VIP*-2 is to the right, where the axis is red, and lesser expression of *VIP*-2^m^ relative to *VIP*-2 is to the left, where the axis is blue. **d,** The allelic imbalance in the AH was significantly greater than in the IN. **e,** Allelic imbalance positively predicted *VIP* expression in the AH, but negatively predicted *VIP* expression in the IN. * p < 0.05

### The degree of allelic imbalance in *VIP* expression differed between AH and IN

Morph differences in *VIP* expression could be explained by differential regulation of the standard (ZAL2) and supergene (ZAL2^m^) alleles (Prichard et al., 2022; Sun et al., 2018). To test for differential expression of the two alleles in AH and IN in heterozygotes, we measured allelic imbalance in WS birds using a multiplex RT-qPCR assay targeting a SNP in the 3’UTR. The ZAL2^m^ allele of *VIP* (*VIP*-2^m^) was overexpressed, relative to the ZAL2 allele (*VIP*-2), in the AH (one-sample t-test: t = 10.95, p < 0.001; Figure 2c) and the IN (one-sample t-test: t = 6.01, p < 0.001; Figure 2c). Comparing the two regions directly showed that the degree of allelic imbalance was significantly greater in the AH than in the IN (F_(1,68)_ = 4.60, p = 0.036; Figure 2d), which demonstrates that the degree to which the two alleles are differentially regulated depends on the region. The local regulatory environment may dictate the extent to which each allele contributes to overall expression; whereas the degree of allelic imbalance in AH positively predicted the overall level of expression in that region (R = 0.367, 95%CI [-0.015, 0.656], p = 0.060; Figure 2e), the association between allelic imbalance and expression in IN was negative (R = −0.409, 95%CI [-0.663, −0.070], p = 0.020; Figure 2e) and the associations in AH and IN were significantly different from each other (Fisher’s Z: p = 0.003).

## Discussion

In this study, we have shown that in a species with a supergene polymorphism, the expression of a single gene predicts two opposing behaviors that form the basis of alternative life history strategies. Together, our findings suggest that the same *cis-*variant of *VIP* produces different functional consequences across hypothalamic regions. Allele-specific expression of *VIP* is differentially regulated across neural circuits that mediate aggression and parental care, consistent with a coordinated behavioral trade-off. Our findings suggest that *cis*-regulatory variation between the alleles works together with region-dependent trans-regulation to bias *VIP* expression in opposite directions across functionally distinct neural circuits, supporting the evolution of alternative life-history strategies in this species.

As was previously reported for a different population of white-throated sparrows (Horton et al., 2020), we found that *VIP* expression differed between the morphs and that the direction of this difference depended on whether the population of VIP neurons is pro-aggression (AH) or pro-parental (IN). The current study replicates and expands these findings, in that we observed a significant morph–by–region interaction: *VIP* expression was elevated in AH but reduced in IN in WS birds relative to TS birds (Figure 2a). Further, expression levels in AH and IN were negatively correlated: birds with relatively high *VIP* expression in AH tended to exhibit low expression in IN, and vice versa (Figure 2b), mirroring the behavioral trade-off. This cross-region anticorrelation indicates a more complex arrangement than two independent effects of morph; rather, it suggests that *VIP* expression across these two regions could be regulated in a coordinated manner.

The expression of *VIP* predicted behavior in directions consistent with a role for this neuromodulator in both aggression and parenting. In the AH, higher *VIP* expression predicted territorial singing (Figure 1c), replicating prior findings (Horton et al., 2020) and further demonstrating the relevance of this population to aggressive behavior (Goodson et al., 2012a, b). In the IN, by contrast, *VIP* expression positively predicted parental provisioning (Figure 1e). These relationships persisted after controlling for morph and sex, indicating that the association between regional *VIP* expression and behavior cannot be explained by morph or sex differences in the behaviors or the gene expression. Thus, expression of the same gene, captured within the supergene, predicts opposing components of the behavioral phenotype.

Our analysis of allele-specific *VIP* expression provides important clues about the role of *cis*-regulatory variation in morph-dependent *VIP* expression. *VIP*-2^m^ was overexpressed in both AH and IN, but more strongly in AH; the difference in the magnitude of bias between the hypothalamic regions suggests that locally distinct regulatory landscapes modulate the effects of *cis*-regulatory variation between the alleles, with morph-specific phenotypic consequences. In AH, as allelic imbalance in favor of *VIP*-2^m^ increased, the total expression also increased (Figure 2e), suggesting that increased total expression in WS birds (Figure 2a) can be attributed to higher transcription activity of *VIP*-2^m^. This finding is consistent with Prichard et al. (2022) who reported lower levels of methylation of *VIP*-2^m^, compared with *VIP*-2, in the hypothalamus.

In the IN, the relationship between allelic imbalance and total *VIP* expression was not as straightforward: as allelic imbalance in favor of *VIP*-2^m^ increased, total transcript abundance declined (Figure 2e). This result could be caused by a morph difference in VIP cell number; if TS birds have more VIP-producing cells in IN than do WS birds, their overall *VIP* expression would be higher even in the face of higher transcription activity on ZAL2^m^ in WS birds. Our previous work has shown, however, that in captive populations, WS birds have more, not fewer, VIP-immunoreactive cells in IN (Maney et al., 2005; Maney et al., 2020). A more likely explanation for the relationship between *VIP*-2^m^ and overall *VIP* expression is that, despite greater *cis*-regulatory activity of the *VIP*-2^m^ allele, total expression depends critically on local trans-regulatory influences (Wittkopp et al., 2004), which typically vary dramatically from tissue to tissue (Pinter et al., 2015). Additionally, increased dominance of *VIP-*2^m^ transcripts in WS birds could co-occur with compensatory or other morph-dependent regulation that reduces overall *VIP* expression even as allelic bias strengthens (Bader et al., 2015). Our findings suggest that the regulatory environment in IN, which houses pro-parental VIP neurons, is distinct from that in AH, where pro-aggression VIP cells reside.

VIP’s role as a mediator of behavioral trade-offs is not a quirk unique to white-throated sparrows. This function of VIP is likely conserved across a wide variety of species. In zebra finches, numbers of immunolabeled VIP neurons and colocalization of VIP with fos protein predicted aggressive and parental behaviors, positively or negatively, depending on the brain region (Kingsbury et al., 2015). Kingsbury and Wilson (2016) argued that VIP is positioned within conserved neural circuits to influence both territoriality and parental care, making it a candidate mechanism for coordinating competing social strategies. We show here that in a species with known allelic differentiation, *VIP* expression is differentially regulated across brain regions in an allele-specific way and allelic imbalance predicts behavioral variation in opposite directions. Thus, rather than merely participating in both behaviors, VIP appears to provide a substrate through which regulatory divergence can partition aggression and parental care, linking a conserved neuropeptide system to alternative life-history strategies.

The trade-off between territorial aggression and parental care is widely hypothesized to be mediated by androgens (reviewed by Hau, 2007; Ketterson & Nolan, 1994; McGlothlin et al., 2007). In the context of our findings, androgen signaling could plausibly influence *VIP* transcription. Morph-based variation in androgen receptor abundance (Grogan et al., 2019) does not map cleanly onto variation in *VIP* expression, however, suggesting that morph differences in *VIP* are not explained simply by the distribution of androgen receptors. Because regulatory variation between *VIP*-2^m^ and *VIP*-2 affects predicted androgen response elements (Prichard, unpublished data), it is possible that androgens have differential effects on allele-specific expression of *VIP*. Testing this hypothesis will require experimental manipulation of androgen signaling combined with allele-specific expression assays.

Dobzhansky (1950) proposed that chromosomal inversions are favored because they capture co-adapted alleles that function optimally together. Evidence from diverse taxa, including fire ants (Purcell et al., 2014), Heliconius butterflies (Joron et al., 2006), and ruffs (Küpper et al., 2016) supports this classic model; chromosomal inversions maintain co-adapted alleles that underlie alternative reproductive or social strategies. In these systems, chromosomal inversions link suites of genes whose combined effects are thought to produce divergent, complex phenotypes encompassing suites of behaviors that are multiply determined (reviewed by Charlesworth, 2016). Our current findings suggest a variation on this theme: competing behaviors might be mediated by regulatory divergence within a single locus that, through pleiotropic effects, itself produces opposing functional consequences.

VIP is unlikely to act alone, however. In white-throated sparrows, aggression and parental behavior are linked also to a gene that neighbors *VIP* inside the supergene: *ESR1*, the gene encoding estrogen receptor alpha. Expression of *ESR1* is elevated in WS birds in the ventromedial arcopallium (AMV), where it predicts aggression (Horton et al., 2014b) and where knockdown eliminates the aggressive phenotype (Merritt et al., 2020). Expression of *ESR1* in a separate region, the medial preoptic nucleus (POM), predicts parental behavior (Horton et al., 2014b). Together, these findings suggest that the inversion polymorphism in this species may coordinate multiple loci with region-specific expression patterns that align with alternative social strategies. Notably, it is not the case that one of the genes affects aggression and the other affects parenting -- *both* genes are involved in *both* behaviors, making it particularly interesting that regulatory variation has arisen separately, within each of these genes, to promote or inhibit the same suite of competing behaviors.

If a single gene, like *ESR1* or *VIP*, can alone mediate multiple behaviors that make up a complex behavioral phenotype, what, then, is the advantage of being captured inside a supergene? The answer may lie in the white-throated sparrow’s mating system. In this species, the morphs mate disassortatively; nearly all breeding pairs comprise one WS and one TS bird (Lowther, 1961; Tuttle et al., 2016). Thus, like the mammalian Y chromosome, the ZAL2^m^ chromosome is typically present in only one member of a breeding pair. The supergene on ZAL2^m^ does not simply bundle together separate behavioral mechanisms; it partitions their effects in ways that generate complex, adaptive phenotypes both within individuals *and across breeding partners.* That this architecture resides on an autosome, rather than a sex chromosome, highlights the extraordinary ways in which inversion polymorphisms can appropriate the logic of sexual differentiation (Charlesworth, 2016).

Our findings demonstrate that when the regulation of a single gene differs across neural circuits, its expression can predict opposing components of a behavioral trade-off. In white-throated sparrows, *VIP* expression is differentially regulated in two hypothalamic cell populations, one associated with aggression and the other with parenting. These patterns are further shaped by allele-specific regulatory variation; the supergene-associated allele contributes to expression in a manner that depends on the local transcriptional environment. Our results suggest a refinement of the classical supergene model. Rather than requiring large sets of co-adapted loci to generate integrated behavioral phenotypes, coordinated trade-offs may arise, at least in part, from individual pleiotropic genes with effects that diverge across tissues. In this framework, chromosomal inversions may facilitate the evolution of alternative strategies not only by linking multiple genes, but also by driving regulatory divergence in individual genes that produces context-dependent, and even opposing, functional outcomes. More broadly, this work highlights how gene regulation across neural circuits can organize complex life-history strategies and provides a mechanistic bridge between genomic architecture and behavioral trade-offs.

## Acknowledgements

This work was supported by NIH Grant 1R01MH082833 to D.L.M. We are grateful to the Boulder Lake Environmental Learning Center for granting us access to their lab space and their property, especially Ryan Hueffmeier, who was immensely helpful with coordination. We would like to thank the staff at the Natural Resources Research Institute in Duluth, MN for providing us with access to a −80 freezer in 2019. This work was made possible thanks to the field technicians Michelle Moyer, Rachel Weisbeck, Edwin Harris, Wren Corvus, Lauren Chronnister, and Emilia Skogen, who helped collect behavioral data, find nests, and collect tissue samples, as well as to Irfan Masud and Reeya Bazari, who reviewed the video footage and scored parental behavior. Lastly, we respectfully acknowledge that this work was conducted on the native, ancestral lands of the Anishinabewaki and Očhéthi Šakówiŋ people.

## Online Methods

### Study Population

All fieldwork was conducted during May – July of 2019 and 2021 on an 18,000 acre property owned by Boulder Lake Environmental Learning Center near Duluth, Minnesota, U.S.A. All research was carried out with the approval of the Emory University Institutional Animal Care and Use Committee and with appropriate local (St. Louis County Land and Minerals Department, Authorization Number A1021002), state (Minnesota Department of Natural Resources, Special Permits 23705 and 30224), and federal (U.S. Geological Survey, USGS Permit Number 23369; U.S. Fish and Wildlife Service permit number MB009702-0) permits.

### Behavior

The individuals in this study comprised two cohorts, referred to here as pre-parental and parental. Territorial behavior was quantified for the pre-parental cohort during the early breeding season, before the start of incubation, and parental behavior was quantified for the parental cohort later in the breeding season when adults were engaged in nestling provisioning. All breeding pairs were heteromorphic (*i.e.*, one TS and one WS adult, for sample sizes refer to Table S1). The birds in the two cohorts were collected from different parts of the property to minimize disturbances and cross-exposure to stimuli (*e.g.*, exposure of parental animals to decoys, see Territorial Behavior, below). Birds were captured for banding an average of 16 days before the first behavioral observation, except for two birds in the parental cohort that were not banded.

All birds were trapped by luring them into mist nets with playback of conspecific song. All birds were fitted with an aluminum USGS identification band and plastic bands in a unique color combination for identification during behavioral observations. We evaluated the breeding stage for each animal in hand by measuring the width and height of the cloacal protuberance, if present, or scoring the development of the brood patch, if present. All birds were then released at the site of capture. We continuously monitored all breeding pairs throughout the breeding season and noted their progression through the breeding stages by observing nesting behaviors such as nest building, incubation, or nestling provisioning. Data collection from the pre-parental cohort ceased once it was determined that most of the females were laying eggs and beginning to incubate. Data collection for the parental cohort began once the nestlings were 4-days post-hatch and continued until the end of July.

#### Pre-parental Cohort

Studies on territorial aggression were conducted during early summer (May 14 – Jun 5, 2019; May 16 – Jun 3, 2021). We determined the boundaries of the territories of breeding pairs on the basis of behavioral observations of the focal pair and neighboring pairs. Territorial aggression was quantified using simulated territorial intrusions (STIs). A caged, male conspecific decoy, captured at least 4.5 miles from the site of the STI, was placed within the territory of a focal pair along with a Bluetooth speaker (model: FBA_AYL-SoundFit) playing conspecific song. All STIs were conducted in the morning (0530-1130 EDT). STIs were performed twice, once on two consecutive days, to present each breeding pair with a male decoy of each morph separately. Decoy morph was balanced across trials (Horton et al., 2012). Each decoy was paired with a unique playback recording (Horton et al., 2014a) and presented only four times: once first and once second for each of the two resident pair types (WSM/TSF, WSF/TSM). Decoys were released near the site of capture.

The STI protocol and the behaviors recorded were as previously described by Horton et al. (2014a). Two observers with unique viewpoints of the decoy and focal bird(s) performed the STIs. After placing the caged decoy and speaker, one of the observers began the playback, and each began their own timer and recorded their observations. If neither member of the focal pair appeared within 15 minutes, the trial was stopped and we returned the next day to try again in a different part of their territory, when possible. Otherwise, when the first member of the focal pair showed interest in the decoy, the observers began a 10-minute countdown. During the following 10-minute period, the observers recorded territorial behavior from each bird in the focal pair. The behaviors that were recorded included singing, latency to first approach, time spent within 5 m of the decoy, time spent within 2 m of the decoy, distance of the closest approach, physical contact with the cage, flights over/toward the cage, other vocalizations (*e.g.*, chips, chip-ups, trills), and copulation solicitations (see Table S2 for more detailed descriptions of the behaviors). At the end of the 10-minute STI, the two observers compared their observations and agreed on the final data set for each trial. Discrepancies were resolved either by using the data from the observer with a better line of sight or by using the average.

#### Parental Cohort

Parental behavior was quantified in the parental cohort during mid-summer (June 15 – July 16, 2019 and June 9 – July 10, 2021) when adults were engaged in nestling provisioning. Video recordings of parental behavior were collected when nestlings at the recorded nest were 5-7 days post-hatch. Nestling age was typically based on the known hatch date, but for nests that were found after hatching, the age of the nestlings was visually evaluated following U.S. Fish & Wildlife guidelines (Jongsomjit et al., 2007). All recordings were made between 0530 – 1200 EDT and during fair weather. Cameras (Sanyo Xacti MPEG-4 AVC/H.264) were placed 2 – 6 ft from the nest, depending on the cover provided by surrounding vegetation. The cameras were left to record for 2-3 hours.

Video recordings were stored on a hard drive until they were evaluated by one or two scorers who were blind to the hypotheses about morph differences. Agreement between scorers for the parental data was tested with the intraclass correlation coefficient (ICC) using icc (*irr* package, Gamer & Lemon, 2019; Koo & Li, 2016). Approximately 28% of the videos were scored by both scorers, and the data from this subset were highly consistent between the scorers (ICC = 0.980 ± 0.018). Behaviors that were quantified from the videos included number of provisioning trips, fecal sac removals, time spent brooding, and time spent at the nest (see Table S3 for more detailed descriptions of the behaviors). The video scorers attributed behavior to each adult parent by observing band colors when possible or plumage coloration (WS or TS) when bands were not visible. All behavior scoring was conducted in BORIS (v. 7.13.9; Friard & Gamba, 2016).

### Tissue Collection

Birds were captured for tissue collection 1-2 days after the second STI for the pre-parental cohort and on post-hatch day 7 or 8 for the parental cohort. Capture by mist net occurred 3.5 ± 5.3 min (mean ± SD; both cohorts combined) after the start of playback. Two blood samples were collected for another study, one from a wing vein directly after capture and the other from the jugular vein, under isoflurane anesthesia, immediately before decapitation (07:45 ± 03:26; MM:SS after capture). Brains were dissected and flash-frozen in powdered dry ice at 03:13 ± 02:02 (MM:SS) min after decapitation. A liver sample for genotyping was collected into a 1.5 ml Eppendorf tube. The brains and liver samples were stored on dry ice for ∼3 hours, then transferred to a −80°C freezer until later transport on dry ice to Emory University. At the time of collection, sex was confirmed by visual inspection of the gonads. For the parental cohort, nestlings were also collected, euthanized by overdose of isoflurane, and used for another study. The age of the nestlings was re-evaluated in hand upon collection (Jongsomjit et al., 2007). If our estimation of the age of the nestlings was adjusted at this stage, then the video from corrected day 6 post-hatch was used for behavioral scoring.

### Microdissection and DNA/RNA extraction

Frozen brains were sectioned at 200µm thickness and thaw-mounted onto microscope slides. Regions containing the AH and IN were microdissected using the Palkovits punch technique (Palkovits, 1985). To target the AH, we took two bilateral punches 1 mm in diameter from two consecutive sections ventral to the anterior commissure (Figure S3). When targeting the IN, we took a single 1mm punch, centered on the midline, from three consecutive sections just rostral to the median eminence (Figure S3). We also collected four 1 mm punches from the hippocampal area of one section (Figure S3) to use for inter-run calibration during qPCR. Punched tissues were preserved in DNA Shield (Zymo; Irvine, CA, USA) and stored at −20°C until RNA extraction. DNA/RNA extractions from brain punches and liver were conducted using the Quick DNA/RNA Microprep Kit (Zymo). Extracted DNA from the brain was stored for use in another study and DNA from the liver was used to confirm both morph and sex by genotyping via PCR. Extracted RNA was quantified by Nanodrop and cDNA converted on the same day with the Transcriptor High Fidelity cDNA Synthesis Kit (Qiagen; Valencia, CA, USA). The cDNA conversions included two negative controls to test for gDNA contamination: a reaction without reverse transcriptase and a no template control.

### *VIP* expression

Expression of the *VIP* gene was measured using RT-qPCR performed using a Roche LightCycler 480 Real-Time PCR System. Two housekeeping genes (HKGs), glyceraldehyde-3-phosphate dehydrogenase (*GAPDH*) and peptidylprolyl isomerase A (*PPIA*), were selected because their expression is stable across individuals and does not differ by morph in brain tissue in this species (Zinzow-Kramer et al., 2014). Primers and probes for *VIP* and HKGs were designed and manufactured by Integrated DNA Technologies (Coralville, Iowa, USA) to target a region that included an intron-exon boundary for each gene (Table S5). Primers were validated with a cDNA template by visualizing PCR amplicons on an agarose gel to confirm specificity in both morphs. Each 10µl qPCR reaction contained 2.5µl of cDNA template diluted 1:10, 5µl of PrimeTime Gene Expression Master Mix (IDT), 1µl of gene-specific primers (10nM), 0.5µl gene-specific probe (5nM), and 1µl Dnase/Rnase free H_2_O. Cycling conditions were 1 cycle at 95°C for 3 minutes, 45 cycles at 95°C for 5 seconds and 60°C for 30 seconds, and 1 cycle at 37°C for 1 minute. Morph and sex were balanced across each qPCR plate, and samples collected in different years were run on separate plates to group the nuisance variables of year and plate together. Each plate included both brain regions (AH and IN) for each individual as well as four inter-run calibrator samples and two negative controls to test for gDNA contamination during cDNA conversion: a negative reverse transcriptase control and a negative template control. The four inter-run calibrator samples were each a unique pool of cDNA extracted from brain tissue collected from the hippocampal region at the same time as the punches for the AH and IN (Figure S3). A standard curve of cDNA with a 1:5 dilution factor was used to assess amplification efficiency. Crossing point / cycle threshold (Cp) values were calculated using the “Abs quant/Second derivative max” method in the LightCycler Program (software v. 1.5.0). All samples were run in triplicate; if the standard deviation of Cp for a triplicate was greater than 0.2 then one technical outlier was removed. If the standard deviation of the sample was still greater than 0.2 after removing a technical outlier, the sample was designated as hyper-variable and either re-run or excluded.

#### Calculating relative expression

Cp values were exported and subsequent analyses were conducted in R (R version 4.3.0) and Rstudio (“Mountain Hydrangea” Release, (583b465e, 2023-06-05) for Windows). The expression of *VIP* relative to HKGs was calculated on the basis of Hellemans et al. (2007). Briefly, Cp values were averaged across technical replicates, and amplification efficiency was calculated from the linear slope of the standard curve (gene-specific efficiency values: *VIP* = 2.033 ± 0.053; *GAPDH* = 2.068 ± 0.058; *PPIA* = 2.026 ± 0.055). The ΔCp for each sample was estimated by standardizing each Cp value to the mean of all samples for that gene/probe. The relative quantity of our gene of interest (*VIP*) was normalized to the geometric mean of the two HKGs.

### *VIP* allelic imbalance

The degree of allelic imbalance in *VIP* expression was quantified using a multiplexed qPCR assay designed and manufactured by Integrated DNA Technologies (Coralville, Iowa, USA). Primers amplified a region of the 3’UTR containing a C/T SNP that was then targeted by allele-specific probes (Table S5). The probe targeting the ZAL2 allele was labeled with 6-FAM fluorescent dye and the probe targeting the ZAL2^m^ allele was labeled with 5Cy5 fluorescent dye. The difference in fluorescence between these two probes allows them to be distinguished in a duplex reaction without color compensation (LightCycler 480 Manual; Roche Diagnostics, Indianapolis, IN). Primers were validated using cDNA from both morphs by visualizing PCR amplicons on an agarose gel to confirm specificity. All samples were run in triplicate on the same plate, and technical outliers were excluded. Only heterozygous WS cDNA samples were used in this assay because TS birds are homozygous for the *VIP*-2 allele and, therefore, do not exhibit allelic imbalance. qPCRs were performed in volumes of 10µl including 5µl of PrimeTime Gene Expression Master Mix (IDT), 1µl of forward and reverse primers (5nM), 0.25µl *VIP*-2^m^-specific probe (10nM), 0.25µl *VIP*-2-specific probe (10nM), and 1µl DNase/RNase free H_2_O. Cycling conditions were 1 cycle at 95°C for 3 minutes, 45 cycles at 95°C for 5 seconds and 60°C for 30 seconds, and 1 cycle at 37°C for 1 minute.

Each plate included a standard curve to confirm that the allele-specific probes had similar and satisfactory amplification of both alleles. This standard curve was made with a 1:5 dilution factor using gDNA samples from heterozygous individuals, which have a 1:1 ratio of the ZAL2:ZAL2^m^ alleles. This standard curve also provided the slope for the amplification efficiency used in subsequent calculations. To confirm that the specificity of the probes was similar across degrees of allelic imbalance, we used gDNA from a rare ZAL2^m^/ZAL2^m^ homozygote (Horton et al., 2013) and a TS bird (*i.e.*, a ZAL2/ZAL2 homozygote) to create a second standard curve of known ratios of ZAL2^m^: ZAL2 gDNA along a linear scale (1:8, 1:4, 1:2, 1:1, 2:1, 4:1, 8:1). The Cps measured from this second standard curve were converted into ratios (ZAL2^m^: ZAL2) and plotted against the expected ratio to test for a linear relationship (Figure S4). In addition to these two standard curves, a sample of gDNA from the ZAL2^m^/ZAL2^m^ homozygote and a TS bird were included on each plate to confirm minimal amplification of the ZAL2 allele in the ZAL2^m^ sample and vice versa. Two negative controls, a negative cDNA reverse transcriptase control and a negative RNA template control were also included to test for gDNA contamination in the cDNA conversion. To adjust for variation between runs, we used four unique cDNA pools created from hippocampal tissue as described above, as calibrators.

#### Calculating allelic imbalance

Cp values were calculated using “Abs quant/Second derivative max” in the LightCycler Program (software v. 1.5.0). The relative expression of each allele, that is, *VIP-*2^m^ / *VIP*-2, was calculated for both the AH and IN for each bird based on methods described by Hellemans et al. (2007) and Merritt et al. (2020). Briefly, the mathematical protocol for this assay was the same as in the relative expression qPCR assay, described above, but instead of *VIP*:(*GAPDH, PPIA*), the relative expression quantified was *VIP*-2^m^:*VIP*-2. Because the primers for this assay were not intron-spanning, gDNA contamination was a concern. Therefore, we tested all samples for gDNA contamination using an intronic qPCR assay. gDNA contamination was calculated as the concentration of intronic signal divided by the concentration of the allele-specific signal. gDNA contamination was minimal for all samples (< 0.007), and no samples were excluded because of gDNA contamination.

### Statistical Analyses

Statistical analyses were conducted in R (version 4.5.1 (2025-06-13 ucrt) -- "Great Square Root") and RStudio ("Apple Blossom" Release (0e924abb, 2026-02-04) for Windows). All data on aggressive behavior were log_10_ transformed, and all data for parental behavior were square-root transformed for statistical analyses. Data on *VIP* expression were log_2_ transformed. For all statistical models that included it, date was defined as the number of days since the last snowfall for each year (*e.g.*, last snowfall on 5/1/19 and data collected on 5/10/19 = 9 days). Skewness/normality of each variable was tested with shapiro.test (stats package; R Core Team). Outliers were identified using grubbs.test (outliers package; Komsta, 2022) and excluded from downstream analyses. All figures were created using the ggplot2 package (Wickham, 2016).

We took a two-step approach to test for differences in behaviors and *VIP* expression between groups, starting with a general linear mixed model (*afex* package; Singmann et al., 2025) and following up with within-group *post-hoc* Tukey HSD tests when appropriate (emmeans package; Lenth, 2023). For example, if we found a significant interaction between morph and sex when testing for a morph difference with Type 3 ANOVA, then we tested for and reported morph differences within sex using Tukey HSD. Specific models are reported with the results in the supplementary material.

To test for relationships between *VIP* expression and behavior independently of morph we conducted partial Pearson’s correlations. We calculated residual values for each variable (*e.g.*, *VIP* expression in AH and total singing) from a general linear mixed model (nlme package; Pinheiro et al., 2023; Pinheiro & Bates, 2000) that controlled for morph, sex, date and qPCR run, which was confounded with year but balanced across morph and sex. Then we used those residual values to test for correlations between *VIP* expression and behavior (stats package; R Core Team). We used Fisher’s Z to compare correlations where appropriate.

We tested for evidence of allelic imbalance using one-sample Student’s t-tests (stats package; R Core Team). To compare allelic imbalance between the two regions, we used a general linear mixed model and a Type 3 ANOVA test (*afex* package; Singmann et al., 2025). We tested for a relationship between *VIP* expression and allelic imbalance using a partial Pearson’s correlation following a similar approach as described above. We estimated residuals for each variable (*e.g.*, *VIP* expression in the AH and allelic imbalance in the AH) from a general linear mixed model (nlme package; Pinheiro et al., 2023; Pinheiro & Bates, 2000) that controlled for sex, date and year. Then we used those residual values to test for correlations between *VIP* expression and allelic imbalance (stats package; R Core Team). We used Fisher’s Z to compare the correlation in AH with that in IN.

## Notes

### Competing Interest Statement

The authors have declared no competing interest.

