## Supplementary Material for "Opposing behavioral roles for a single gene in a species with a supergene polymorphism"

### Supplementary Tables

*Table S1.* Sample sizes within year, breeding stage, morph, and sex.

| Sample Sizes (n = 74) | | | | |
|---|---|---|---|---|
| **Year** | **Breeding Stage** | **Morph** | **Sex** | **n** |
| 2019 (n = 35) | Pre-parental (n = 20) | WS | M | 5 |
| | | | F | 5 |
| | | TS | M | 5 |
| | | | F | 5 |
| | Parental (n = 15) | WS | M | 3 |
| | | | F | 5 |
| | | TS | M | 4 |
| | | | F | 3 |
| 2021 (n = 39) | Pre-parental (n = 18) | WS | M | 6 |
| | | | F | 3 |
| | | TS | M | 6 |
| | | | F | 3 |
| | Parental (n = 21) | WS | M | 6 |
| | | | F | 4 |
| | | TS | M | 5 |
| | | | F | 6 |

*Table S2.* Morph differences in aggressive/territorial behaviors.

**Total Singing**

| Description | Fixed Effect | DF | F-value | p-value | |
|---|---|---|---|---|---|
| The total number of songs produced by the focal animal | Morph | 39 | 66.6 | <0.001 | * |
| | Sex | 39 | 23.7 | <0.001 | * |
| | Morph:Sex | 40 | 0.20 | 0.656 | |

**Latency to Respond to Decoy**

| Description | Fixed Effect | DF | F-value | p-value |
|---|---|---|---|---|
| Time from start of playback until the focal animal showed interest in the decoy | Morph | 40 | 0.600 | 0.444 |
| | Sex | 40 | 0.63 | 0.431 |
| | Morph:Sex | 40 | 1.020 | 0.318 |

**Time w/in 5m**

| Description | Fixed Effect | DF | F-value | p-value | |
|---|---|---|---|---|---|
| Duration of time the focal animal spent within a 5m radius of the decoy | Morph | 40 | 0.41 | 0.525 | |
| | Sex | 40 | 4.54 | 0.039 | * |
| | Morph:Sex | 40 | 0.03 | 0.864 | |

**Time w/in 2m**

| Description | Fixed Effect | DF | F-value | p-value |
|---|---|---|---|---|
| Duration of time the focal animal spent within a 2m radius of the decoy | Morph | 40 | 0.68 | 0.413 |
| | Sex | 40 | 0.85 | 0.362 |
| | Morph:Sex | 40 | 0.12 | 0.728 |

**Closest Proximity to Decoy**

| Description | Fixed Effect | DF | F-value | p-value |
|---|---|---|---|---|
| The approximate distance in meters of the closest approach to the decoy | Morph | 40 | 2.84 | 0.100 |
| | Sex | 40 | 1.15 | 0.291 |
| | Morph:Sex | 40 | 0.38 | 0.540 |

**Total Flights**

| Description | Fixed Effect | DF | F-value | p-value | |
|---|---|---|---|---|---|
| Total number of flights over, toward, and around the decoy | Morph | 40 | 8.27 | 0.006 | * |
| | Sex | 40 | 5.82 | 0.020 | * |
| | Morph:Sex | 40 | 2.34 | 0.134 | |

**Trills**

| Description | Fixed Effect | DF | F-value | p-value | |
|---|---|---|---|---|---|
| A vocalization exhibited during copulation solicitation (typically female) and aggressive encounters (typically male) | Morph | 40 | 8.89 | 0.005 | * |
| | Sex | 40 | 42.3 | <0.001 | * |
| | Morph:Sex | 40 | 7.78 | 0.008 | * |
| | *Post-hoc* tests for morph differences | | | | |
| | **Sex** | **n** | **t-value** | **p-value** | |
| | Females | 14 | -3.655 | <0.001 | * |
| | Males | 21 | -0.136 | 0.893 | |

**Other Vocalizations**

| Description | Fixed Effect | DF | F-value | p-value |
|---|---|---|---|---|
| Chips, chip-ups, pinks, etc. | Morph | 40 | 0.09 | 0.761 |
| | Sex | 40 | 0.11 | 0.744 |
| | Morph:Sex | 40 | 0.86 | 0.359 |

A significant main effect of morph indicates that WS > TS. A significant main effect of sex indicates M > F except for trills which F > M. In *post-hoc* tests for morph differences, a negative t-value indicates tan-striped (TS) < white-striped (WS), and a positive t-value indicates WS < TS.

Statistical model: $\log_{10}$(behavior + 1) ~ morph * sex + date + 1|year

*Table S3.* Morph differences in parental behaviors.

**Provisioning Rate**

| Description | Fixed Effect | DF | F-value | p-value | |
|---|---|---|---|---|---|
| Total number of provisioning trips to the nest per hour of video | Morph | 34 | 0.79 | 0.379 | |
| | Sex | 34 | 3.39 | 0.074 | |
| | Morph:Sex | 34 | 5.36 | 0.027 | * |

*Post-hoc* morph differences

| Sex | n | t-value | p-value |
|---|---|---|---|
| Females | 21 | -1.077 | 0.290 |
| Males | 18 | 2.166 | 0.038 |

**Latency to Return to Nest**

| Description | Fixed Effect | DF | F-value | p-value | |
|---|---|---|---|---|---|
| Duration (sec) before the focal animal returned to the nest after the disturbance of setting up the camera | Morph | 33 | 2.82 | 0.103 | |
| | Sex | 33 | 3.43 | 0.073 | |
| | Morph:Sex | 34 | 5.71 | .0.023 | * |

*Post-hoc* morph differences

| Sex | n | t-value | p-value | |
|---|---|---|---|---|
| Females | 21 | 0.581 | 0.0565 | |
| Males | 18 | -2.743 | 0.010 | * |

**Total Visits /hr**

| Description | Fixed Effect | DF | F-value | p-value | |
|---|---|---|---|---|---|
| Total number of trips to the nest, either with or without food for the nestlings, per hour of video | Morph | 34 | 0.38 | 0.544 | |
| | Sex | 34 | 4.30 | 0.046 | * |
| | Morph:Sex | 34 | 4.00 | 0.054 | |

**Proportion of Time Spent at Nest**

| Description | Fixed Effect | DF | F-value | p-value | |
|---|---|---|---|---|---|
| Total proportion of time spent at the nest, regardless of activity (e.g., brooding, feeding nestlings, etc) | Morph | 34 | 0.04 | 0.836 | |
| | Sex | 34 | 19.43 | <0.001 | * |
| | Morph:Sex | 34 | 0.01 | 0.717 | |

**Proportion of Time Spent Brooding**

| Description | Fixed Effect | DF | F-value | p-value | |
|---|---|---|---|---|---|
| Total proportion of time spent brooding (i.e., making physical contact with nestlings/brood patch) | Morph | 34 | 0.3 | 0.586 | |
| | Sex | 34 | 13.86 | <0.001 | * |
| | Morph:Sex | 34 | 0 | 0.958 | |

**Fecal Sac Removals**

| Description | Fixed Effect | DF | F-value | p-value | |
|---|---|---|---|---|---|
| Total count of the number of fecal sacs removed from the nest per hour of video | Morph | 34 | 3.41 | 0.073 | |
| | Sex | 34 | 0.04 | 0.846 | |
| | Morph:Sex | 34 | 4.44 | 0.043 | * |

*Post-hoc* morph differences

| Sex | n | t-value | p-value | |
|---|---|---|---|---|
| Females | 21 | -0.235 | 0.816 | |
| Males | 18 | 2.655 | 0.012 | * |

A significant main effect of sex means F > M. In *post-hoc* tests for morph differences, a negative t-value indicates tan-striped (TS) < white-striped (WS), and a positive t-value indicates WS < TS.
Statistical model: sqrt(provisioning trips) ~ morph * sex + date + 1|year

*Table S4.* Main effects of morph, sex, and region on *VIP* expression and their interactions.

| Fixed Effect | DF | F-value | p-value | |
|---|---|---|---|---|
| Morph | 119 | 0.45 | 0.503 | |
| Sex | 119 | 3.8 | 0.054 | |
| Region | 119 | 68.87 | <0.001 | * |
| Morph:Sex | 119 | 1.38 | 0.242 | |
| Morph:Region | 119 | 12.92 | <0.001 | * |
| Sex:Region | 119 | 11.42 | <0.001 | * |
| Morph:Sex:Region | 119 | 1.84 | 0.178 | |

Statistical model: log2(RQR.VIP) ~ Morph * Sex * Region + (1|SampleName) + (1|SampleName/qPCR run)

*Table S5.* Nucleotide sequences of primers and probes for RT-qPCR assays.

**Vasoactive Intestinal Peptide qPCR Assay**

| Gene | Primer Forward | Primer Reverse | Probe |
|---|---|---|---|
| VIP | AAGAAGCCAGGAAGAGCTAAAT | CTTTACCAGGTGTCCTTCAGAG | TTCTGTAGATGAGCTGCTGAGCCA |
| GAPDH | CATCACAGCCACACAGAAGA | CTCCAGTAGATGCTGGGATAATG | CTTCGGCATTGTGGAGGGTCTCAT |
| PPIA | GTGCCGAAGACAGCAGAAA | GCACATGAACCCAGGAATGA | TAGCCAAATCCCTTCTCACCAGTGC |

**Vasoactive Intestinal Peptide Allele-Specific qPCR Assay**

| Gene | Primer Forward | Primer Reverse | Probe |
|---|---|---|---|
| $\text{VIP}^{2}$ | ATGTTGTGAAGTGAAAGTTGTG | GCTTTCAGTAGAGAATGCTAGA | TC+A+G+CTT+TTAT+T+A+CG |
| $\text{VIP}^{2m}$ | ATGTTGTGAAGTGAAAGTTGTG | GCTTTCAGTAGAGAATGCTAGA | ATC+A+A+CT+TT+T+ATTA+C+GT |

### Supplementary Figures

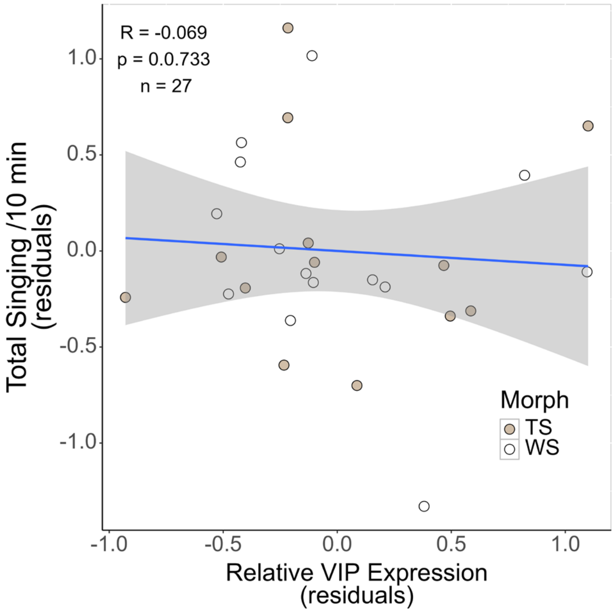

***Figure S1.*** **Correlation between expression of vasoactive intestinal peptide (*VIP*) in the IN and territorial singing.** In contrast to expression in the anterior hypothalamus (Fig. 2c), expression in the infundibular nucleus (IN, x-axis) was not correlated with territorial singing (y-axis).

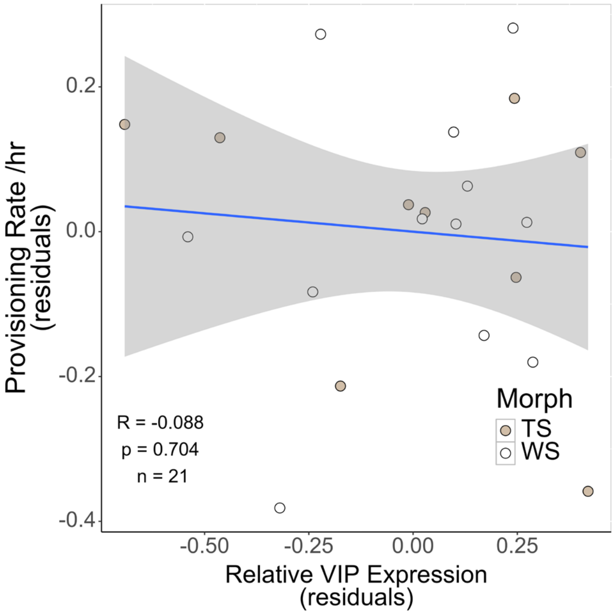

***Figure S2.*** **Correlation between vasoactive intestinal peptide (*VIP)* expression in the AH and nestling provisioning.** In contrast to expression in the infundibular nucleus (Fig. 1e), *VIP* expression in the anterior hypothalamus (AH, x-axis) was not correlated with nestling provisioning (y-axis).

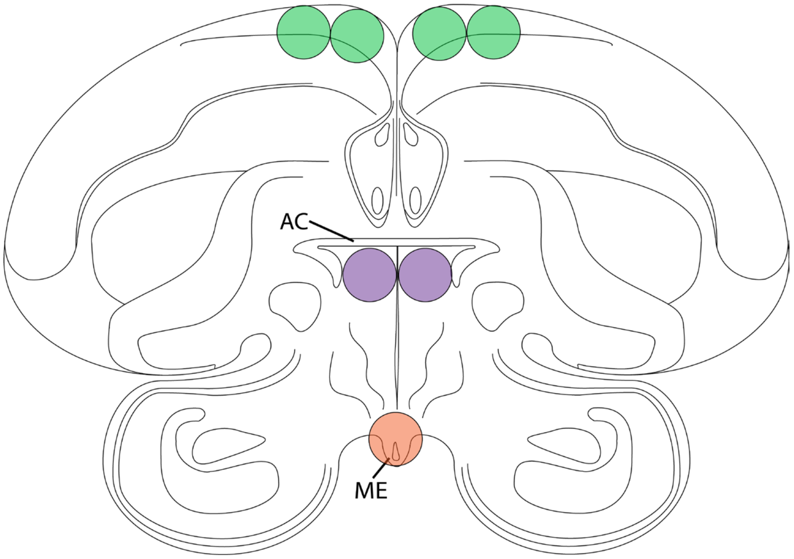

***Figure S3.*** **Anatomical locations of punches of the anterior hypothalamus (AH), infundibular nucleus (IN), and hippocampal area.** The AH was located using the anterior commissure (AC) as a landmark and bilateral punches 1mm in diameter (purple circles) were taken from two consecutive 200µm sections. The IN was located using the median eminence (ME) as a landmark. Punches 1mm in diameter, centered on the midline (orange circle), were taken from three consecutive 200µm sections. Punches of the hippocampal area (green circles, 1mm diameter) were also collected and used for inter-run calibration during qPCR. Illustration by M. R. Prichard.

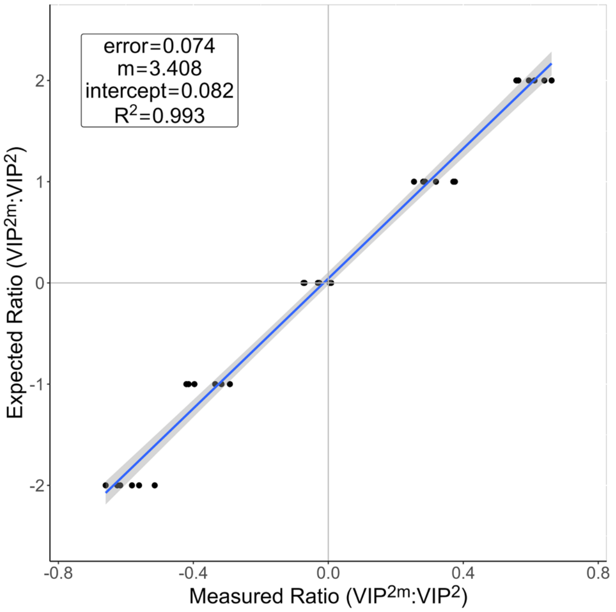

***Figure S4.*** **Standard curve used for the assay of allelic imbalance ($ZAL2^m$: ZAL2).** The standard curve was calculated from samples of gDNA from ZAL2/ZAL2 and $ZAL2^m/ZAL2^m$ homozygotes, combined at known ratios (1:8, 1:4, 1:2, 1:1, 2:1, 4:1, 8:1). The x-axis indicates the ratios calculated from the samples in the standard curve and the y-axis indicates the expected ratio for each sample. The intercept and the $R^2$ suggest that the specificities of the two allele-specific probes were approximately equal across a range of allelic imbalance ratios.
